# Validation of prostate cancer risk variants by CRISPR/Cas9 mediated genome editing

**DOI:** 10.1101/337022

**Authors:** Xing Wang, James E. Hayes, Xing Xu, Xiaoni Gao, Dipti Mehta, Hans G. Lilja, Robert J. Klein

## Abstract

GWAS have identified numerous SNPs associated with prostate cancer risk. One such SNP is rs10993994. It is located in the *MSMB* promoter, associates with *MSMB* encoded β-microseminoprotein prostate secretion levels, and is associated with mRNA expression changes in *MSMB* and the adjacent gene *NCOA4.* In addition, our previous work showed a second SNP, rs7098889, is in LD with rs10993994 and associated with *MSMB* expression independent of rs10993994. Here, we generate a series of clones with single alleles removed by double guide RNA (gRNA) mediated CRISPR/Cas9 deletions, through which we demonstrate that each of these SNPs independently and greatly alters *MSMB* expression in an allele-specific manner. We further show that these SNPs have no substantial effect on the expression of *NCOA4.* These data demonstrate that a single SNP can have a large effect on gene expression and illustrate the importance of functional validation to deconvolute observed correlations. The method we have developed is generally applicable to test any SNP for which a relevant heterozygous cell line is available.

**Author summary:** In pursuing the underlying biological mechanism of prostate cancer pathogenesis, scientists utilized the existence of common single nucleotide polymorphisms (SNPs) in human genome as genetic markers to perform large scale genome wide association studies (GWAS) and have so far identified more than a hundred prostate cancer risk variants. Such variants provide an unbiased and systematic new venue to study the disease mechanism, and the next big challenge is to translate these genetic associations to the causal role of altered gene function in oncogenesis. The majority of these variants are waiting to be studied and lots of them may act in oncogenesis through gene expression regulation. To prove the concept, we took rs10993994 and its linked rs7098889 as an example and engineered single cell clones by allelic-specific CRISPR/Cas9 deletion to separate the effect of each allele. We observed that a single nucleotide difference would lead to surprisingly high level of *MSMB* gene expression change in a gene specific and tissue specific manner. Our study strongly supports the notion that differential level of gene expression caused by risk variants and their associated genetic locus play a major role in oncogenesis and also highlights the importance of studying the function of *MSMB* encoded β-MSP in prostate cancer pathogenesis.

## Introduction

Though genome wide association studies (GWAS) have identified numerous genetic polymorphisms associated with increased risk of disease, the function of these variants remains largely unknown. Just as 99% of the human genome is non-coding sequence, the vast majority of GWAS identified risk associated single nucleotide polymorphisms (SNPs), and SNPs with which they are in high linkage disequilibrium (LD), are located within the non-coding region of the genome [1–3]. They are enriched in regulatory regions of the genome and may alter regulatory elements and expression of nearby genes based on studies on epigenetics and expression quantitative trait loci (eQTLs)[2–7]. Consistent with this, SNPs in regulatory regions appear to be under evolutionary constraint[8]. Experimental validation of this hypothesis has been restricted to a few targeted examples[9–19]. A recent loss-of-function screen in 501 cancer cell lines revealed that 82% of cancers depend on RNA expression level[20], far exceeding mutations and copy number changes, underscoring the importance to study the role of cancer risk variants in gene expression.

Prostate cancer, the second leading cause of cancer-related death in men in the United States, is a good model to investigate the mechanism through which common variants influence disease risk. Prostate cancer is highly heritable, as evidenced by increased rates of comorbidity among monozygotic twins compared to dizygotic twins[21–23]. To date, approximately 100 independent risk SNPs have been identified [24]. These variants tend to be associated with gene expression changes in normal prostate tissue, prostate cancer, and prostatic secretions [25–30] and are found more often in prostatic regulatory regions[31,32]. For instance, the prostate cancer risk SNP rs10993994 is located in the promoter of *MSMB*, which encodes β-microseminoprotein (β-MSP), one of the most abundant proteins in prostate secretions. This SNP is associated with changes in the mRNA level of *MSMB* as well as the nearby androgen receptor co-regulator NCOA4[25,30,33]. Levels of β-MSP in both blood and semen are also associated with this SNP, as are levels of the prostate-secreted proteins prostate-specific antigen (PSA; gene name *KLK3*) and human kallikrein-related peptidase 2 (hK2; gene name *KLK2*)[30]. Rs10993994 is in a 33kb LD block, within which we had also found a second SNP, rs7098889, that is associated with β-MSP levels in prostate tissue independent of rs10993994[30].

With the advent of genome editing tools in mammalian cells such as CRISPR/Cas9[34], it is possible to envision testing a large number of loci for their effect on target gene expression and phenotype. Here, we have chosen to use the highly efficient paired gRNA system[34,35] to delete a candidate regulatory region. This way, in a heterozygote cell line, we can disentangle the effect of each of the two alleles. Here, we apply this system to rs10993994 and rs7098889. We demonstrate that each of these SNPs independently and greatly alters *MSMB* expression in an allele-specific manner. We further show that these SNPs have no substantial effect on the expression of *NCOA4*, nor do they have a direct effect on the prostate secreted proteins hK2 and PSA. These data demonstrate that a single SNP can have a large effect on gene expression and illustrate the importance of functional validation to deconvolute observed correlations. The method we have developed is generally applicable to test any SNP for which a relevant heterozygous cell line is available.

## Results

### Rs7098889 is found in the most prostate-specific enhancer

To determine the causal allele(s) for prostate cancer associated risk SNPs we hypothesized that causal variants would likely be present in prostate specific regulatory elements. Using data on active enhancers from the FANTOM5 consortium analysis of enhancer RNAs[5] (http://slidebase.binf.ku.dk/humanenhancers/selector), we found that the most prostate specific enhancer is located 5kb upstream of the *MSMB* promoter and overlaps rs7098889 (Table 1; Fig 1).

**Table 1.** Top 10 prostate specific enhancers defined by eRNA expression in the FANTOM5 project. Rs7098889 is positioned at chr10:51544475.

| Chr | Position (build 37) | Prostate % of Expression | Tags Per Million |
| --- | --- | --- | --- |
| 10 | 51544405-51544819 | 95.43% | 0.078 |
| 6 | 151709507-151709685 | 87.62% | 0.157 |
| 1 | 59278592-59278937 | 85.02% | 0.078 |
| 3 | 131961835-131962144 | 78.61% | 0.078 |
| 20 | 44632907-44633142 | 76.06% | 0.078 |
| 3 | 57960548-57960842 | 75.56% | 7.221 |
| 22 | 30654777-30654885 | 75.15% | 0.078 |
| 1 | 41915297-41915638 | 71.56% | 0.549 |
| 8 | 18523718-18524110 | 69.58% | 0.942 |
| 16 | 1150049-1150415 | 67.04% | 0.078 |

**Fig 1.**
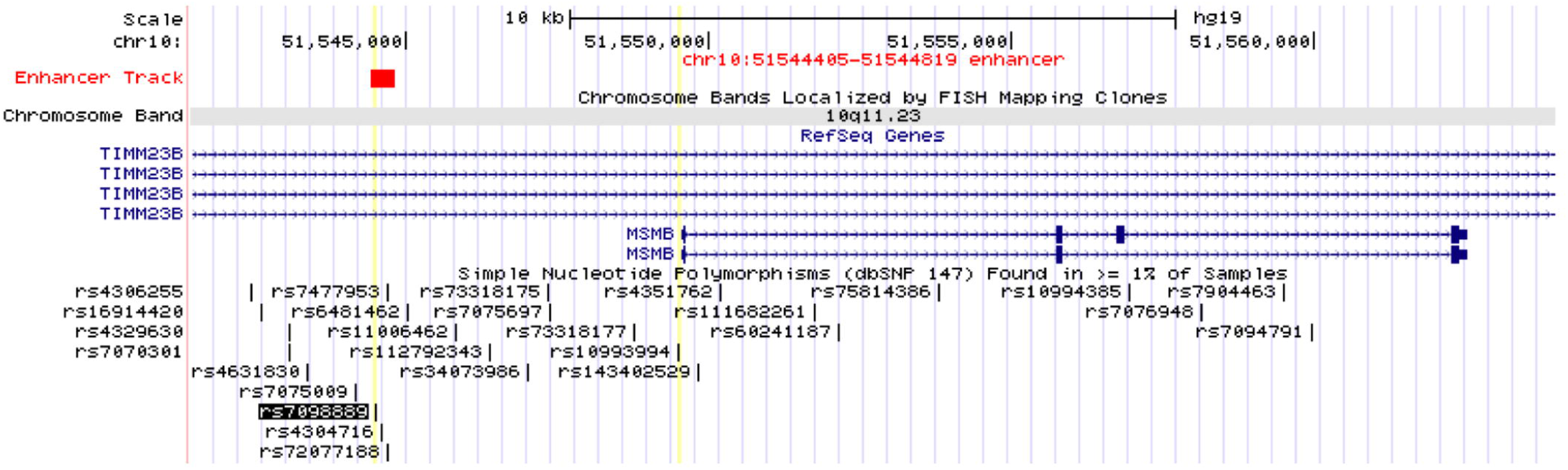
Position of prostate-specific enhancer chr10:51544405-51544819 defined by FANTOM5 project.

### CRISPR/Cas9 mediated deletion of the region flanking rs7098889 leads to significant increase of MSMB expression

We therefore used paired gRNA mediated CRISPR/Cas9 deletion to remove the 191bp region flanking rs7098889 in two heterozygous cell lines -- LNCaP (prostate cancer) and AGS (gastric cancer) (Fig 2A and Fig 2B). Efficient deletion of the region was confirmed by genomic PCR (Fig 2B), though we noted another variant (rs4304716, r^2^=0.87 with rs7098889) in this deletion region as well. We next examined gene expression levels in these bulk transfected cells and found that *MSMB* exhibited 9.5-fold overexpression after deletion of this regulatory region in LNCaP, but not in AGS (Fig 2C). In contrast, *NCOA4*, whose expression is associated with rs10993994 and rs7098889 in GTEx eQTL data[7], was not significantly altered after deletion. Similarly, neither *KLK2* (codes for hK2) nor *KLK3* (codes for PSA) levels were altered after deleting this element (S1 and S2 Figs). The over-expressed transcripts were translated into β-MSP protein and secreted from LNCaP (Fig 2D and Fig 2E). Similar MSMB transcript overexpression was also observed in bulk transfected RWPE-1, another prostate cancer cell line (S3 Fig). These data suggest that rs7098889 is located at a regulatory region that strongly and specifically regulates *MSMB* expression in prostate tissue.

**Fig 2.**
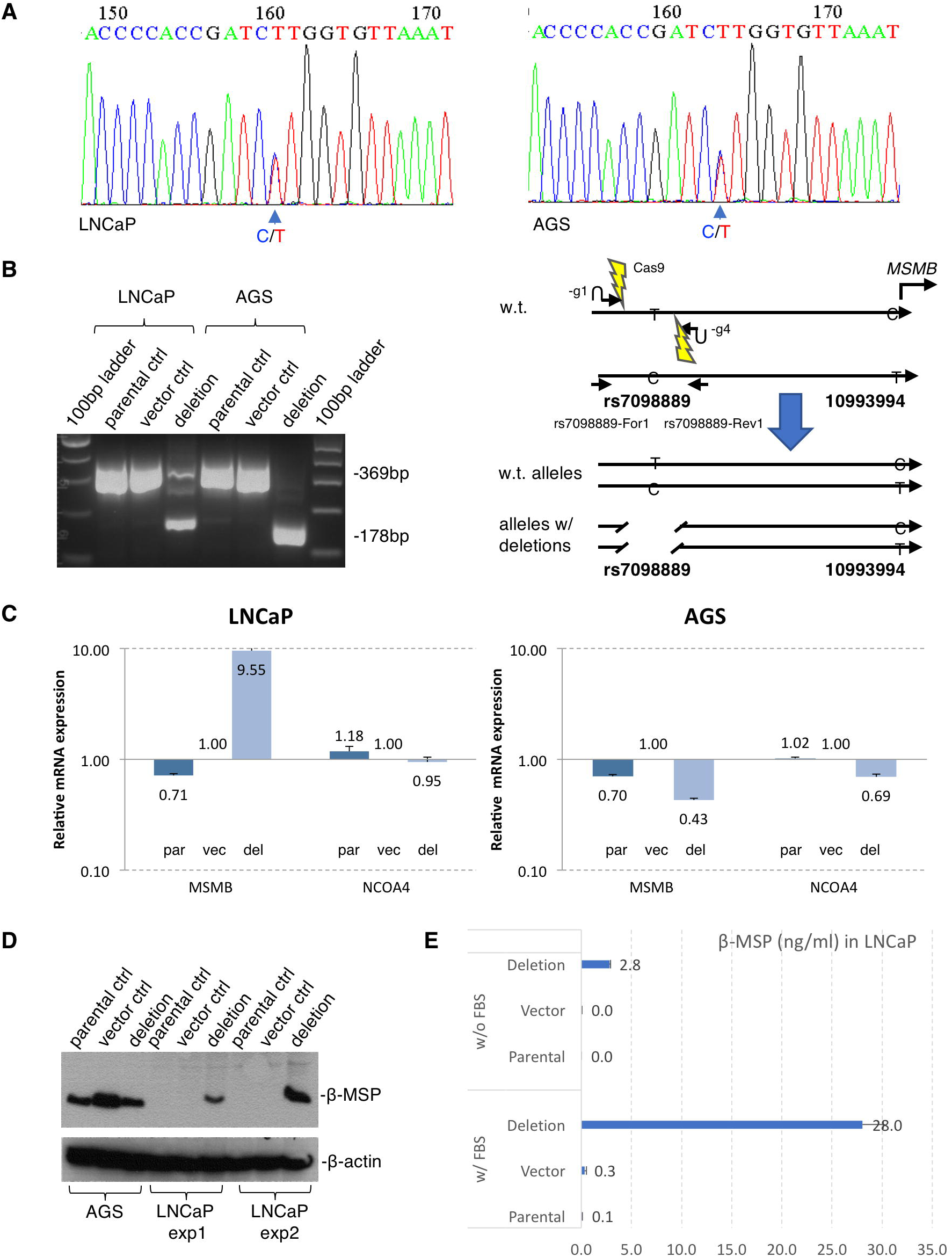
CRISPR/Cas9 mediated deletion of 191bp region flanking rs7098889 confers significant increase of *MSMB* expression. (A)Sanger sequencing showed both LNCaP and AGS cells are heterozygous (C/T) at rs7098889 site.(B)CRISPR/Cas9 mediated rs7098889 deletion was created by paired guide RNAs (rs7098889-g1 and -g4) transfection followed by puromycin selection. The deletion was confirmed by PCR amplification with primer pair (rs7098889-For1 and -Rev1) flanking the deleted region, PCR product runs at 178bp on agarose gel with deletion, and at 369bp without deletion. (C) Real time qPCR showed 9.5 folds *MSMB* over-expression in prostate cancer LNCaP cells with bulk transfection but not the gastric cancer AGS cells. The expression of downstream *NCOA4* gene is barely affected. (D) Western blot showed that the *MSMB* protein product β-MSP is significantly up-regulated in LNCaP cells with deletion, but not in AGS cells. (E) ELISA assay showed that the secreting β-MSP level significantly up-regulated in LNCaP cells with deletion either in the presence (28.0 ng/ml) or absence (2.8ng/ml) of FBS in cell culture.

### Rs7098889 T allele, but not the C allele, confers about 300-fold increase of *MSMB* expression

We next generated single-cell clones from the transfected cells and identified derivative lines that have each of the possible genotypes - del/del, del/T, and del/C - by titrating the transfected plasmids expressing gRNAs/Cas9 to reduce efficiency and increase the chance of getting heterozygous clones (Fig 3A and Fig 2B). Again, in the bulk transfected cells there is a dramatic increase in *MSMB* expression (Fig 3C). Sub-cloning of these cells followed by Sanger sequencing (S1 File) confirmed three T clones (T/del for clone 3, 4, 5), two C clones (C/del for clone 6, 7), as well as two clones with homozygous deletion (del/del for clone 1, 2). Since LNCaP is aneuploid, we also included the upper bands for genotyping to ensure no additional copies of the intended target allele remained. Surprisingly, for every single clone we analyzed, the remaining alleles were always identical (either T or C) at locus rs7098889 (S1 File).

**Fig 3.**
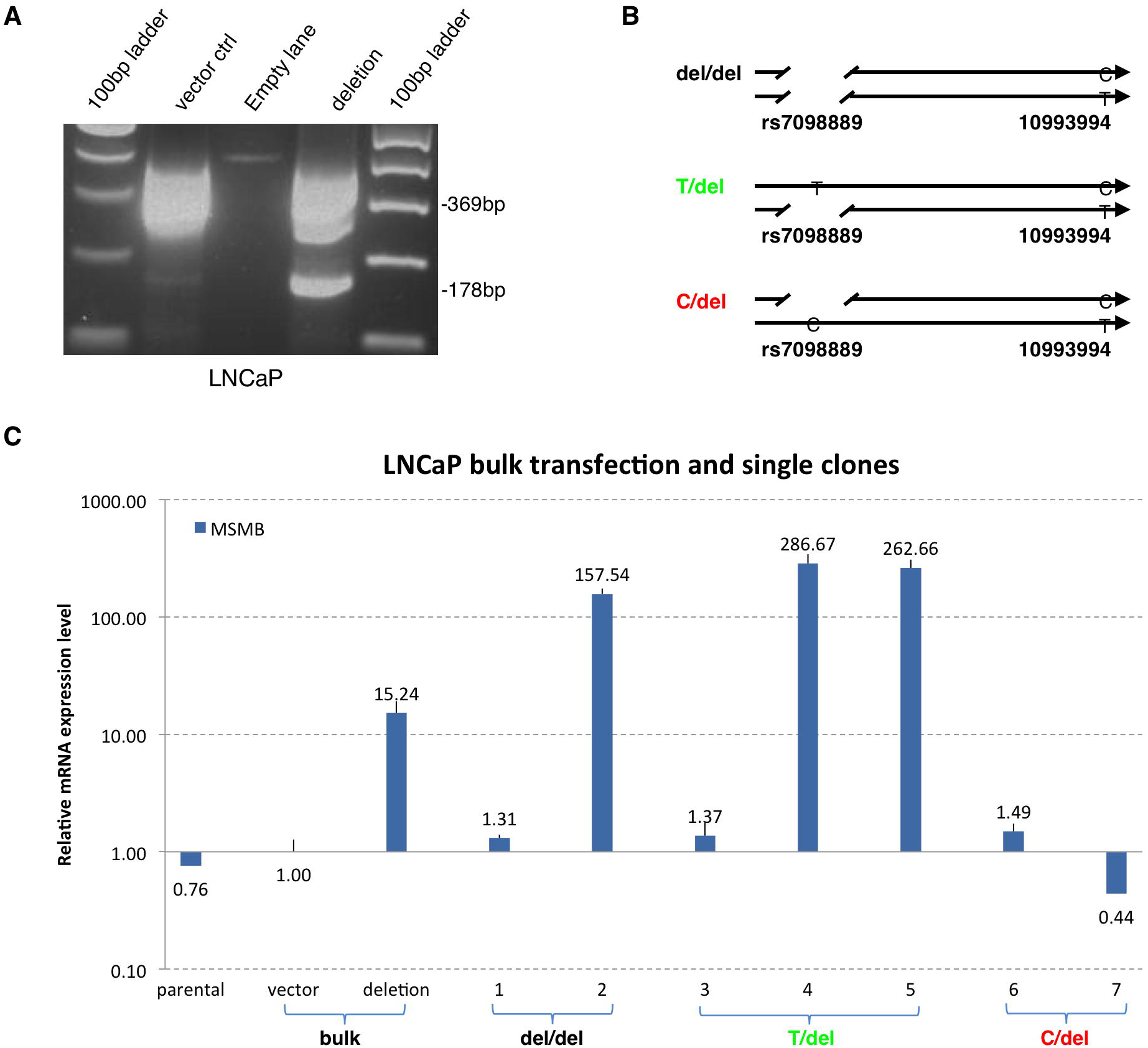
Single clone screening of LNCaP cells with rs7098889 deletion reveals allelic and dramatic *MSMB* over-expression. (A) Transfection titration generated LNCaP bulk cells with lower deletion efficiency for better isolating heterozygous single clones. (B) Illustration of single clone genotypes with homozygous (del/del) and heterozygous deletion (T/del and C/del). (C) Real time qPCR showed dramatic MSMB over-expression (262 and 286 folds) in two (clone 4, 5) out of three clones with rs7098889 T allele (clone 3, 4,5) but not the C allele (clone 6, 7). Bulk deletion with lower deletion efficiency, thus more heterozygous alleles, generates 15 folds over-expression (lane 3) compared to 9.55 folds from previous experiment (Fig. 1C).

Two out of three of the T clones (clones 4, 5) express MSMB close to 300 fold higher than baseline (Fig 3C), while no overexpression is observed in the C clones (clones 6, 7). In the case of the double deletion, one of the two clones overexpressed *MSMB*. To the best of our knowledge, this is the first example of a GWAS-identified risk SNP that unequivocally shows such dramatic effect on gene expression in a highly tissue specific manner. Notably, this effect is very specific to *MSMB* expression alone, as no significant change in adjacent *NCOA4* expression is observed, nor is a trans effect on expression of *KLK2* or *KLK3* present (S2 Fig).

### Allele specific expression of *MSMB* in LNCaP cells

A single nucleotide variant (SNV) 360A/T unique to and heterozygous in LNCaP was identified by Sanger sequencing in the last exon of *MSMB.* We used this SNV as a marker to trace allelic origin of the transcripts. Overexpressed *MSMB* transcripts from all *MSMB* high expression clones (clones 2, 4, 5) and the bulk LNCaP cells all came from the 360T allele (Fig 4B). The control LNCaP parental cells and the empty vector control only express very low basal level of *MSMB.* In these cases, the transcripts come from both alleles (Fig 4B, 360A/T)

**Fig 4.**
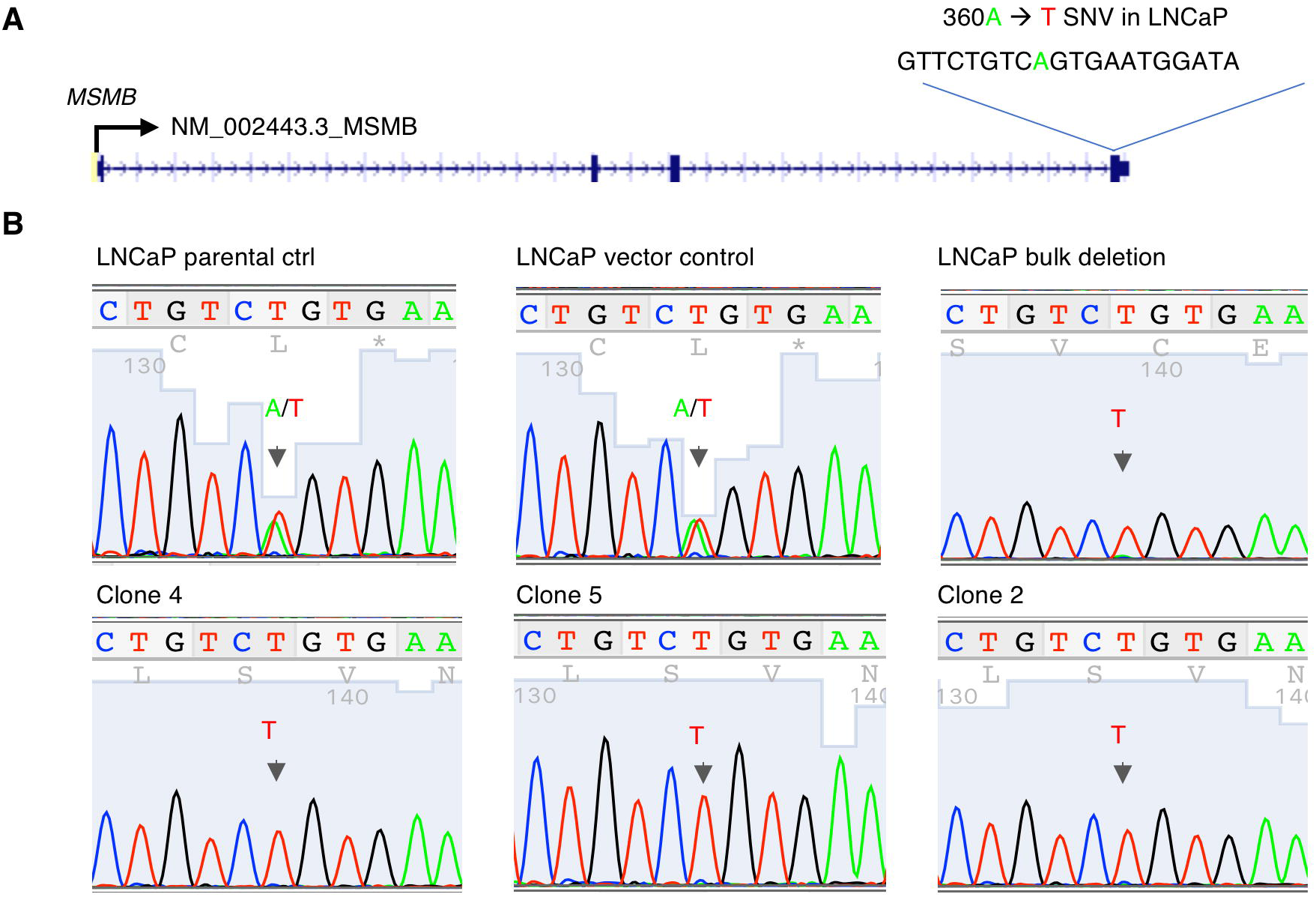
Allelic expression of *MSMB* in LNCaP cells. A)Demonstration of 360A/T single nucleotide variant (SNV) located in the last exon of *MSMB* gene. (B)Transcripts from all the MSMB high expressing clones (clone 2, 4, 5) and the LNCaP bulk deletion came from the 360T allele examined by PCR followed by Sanger sequencing of the last exon of MSMB gene flanking the 360A/T heterozygous site.

### Rsl0993994 C allele, but not the T risk allele, also confers significant *MSMB* over-expression

Previous work at this locus suggested that rs10993994 may be the causal SNP for prostate cancer risk as it localizes in the promoter region of *MSMB* and alters the ability of the promoter to drive expression in a reporter assay[33,37]. Thus, we performed a similar CRISPR/Cas9 mediated genome editing experiment to validate its role in the regulation of *MSMB* gene expression (Fig 5A). We deleted a 205bp region flanking rs10993994. In the bulk transfected LNCaP cells, this deletion results in 2.8 fold overexpression of *MSMB* (Fig 5B). No overexpression was observed either in the gastric cancer AGS cell line or of the immediate downstream NCOA4 gene (Fig 5B). Single cell clones were then generated from the bulk transfected LNCaP cells and resulted in three heterozygous clones with the C allele deletion (T/del for clone 2, 3, 4); three heterozygous clones with T allele deletion (C/del for clone 5, 6, 7); and one clone with homozygous deletion (del/del for clone 1). Even though the deletion removed the majority of the *MSMB* promoter including the TATA box (S2 File), significant overexpression of *MSMB* was observed in two of the three C/del clones (Fig 5C), potentially by transcription from an alternate promoter. Such overexpression was not observed in any of the T/del or del/del clones. Also, we did not observe substantial change of *NCOA4, KLK3* and *KLK2* gene expression in any of these clones (S4 Fig). The result suggests the existence of a strong transcription repression mechanism mediated through both the rs7098889 and the rs10993994 loci. Furthermore, the fact that both loci have dramatic effects on *MSMB* expression in prostate tissue supports the hypothesis that the association of these SNPs with prostate cancer risk is mediated through their regulation of *MSMB* gene expression.

**Fig 5.**
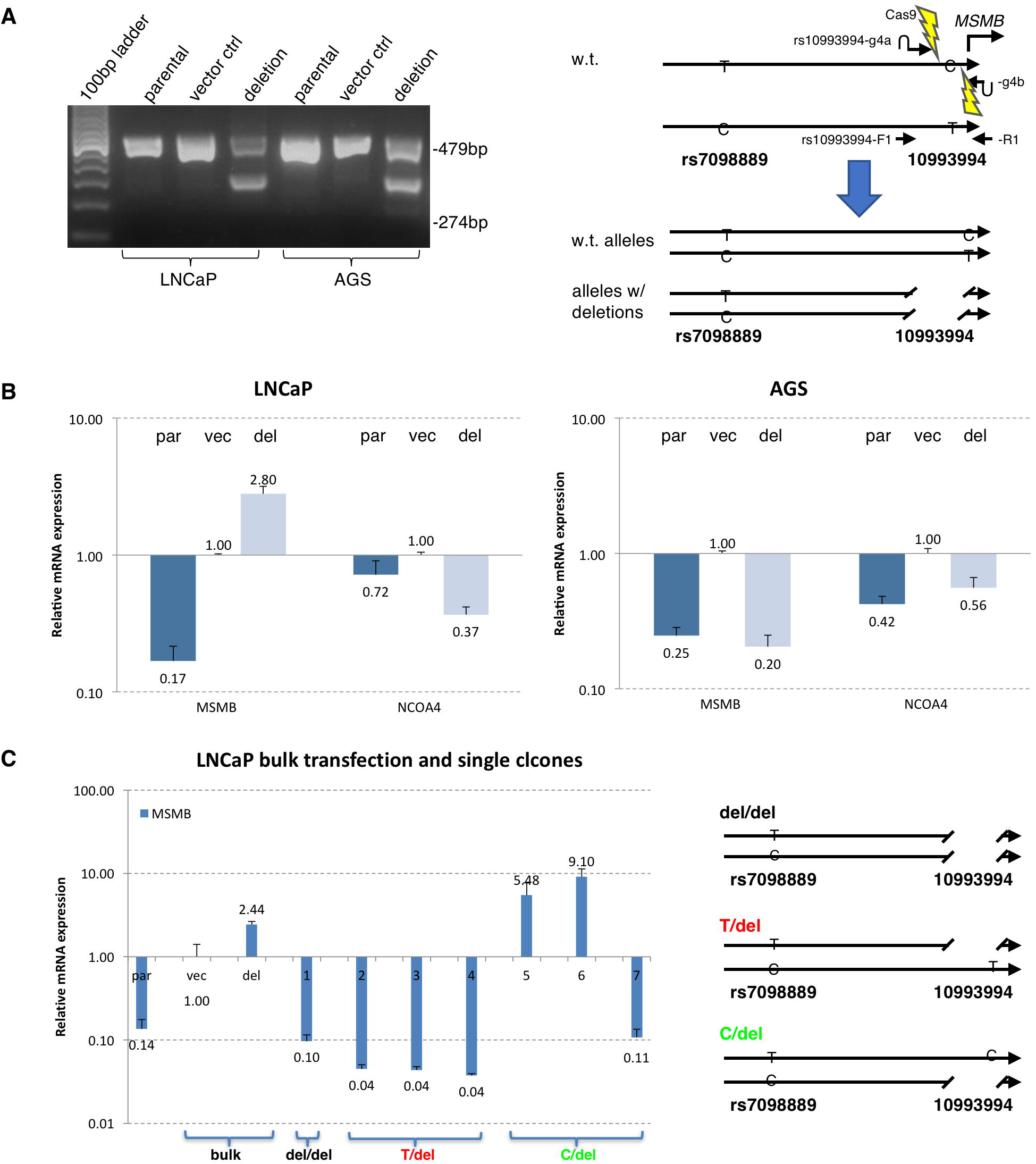
CRISPR/Cas9 mediated deletion of 205bp rs10993994 flanking region confers significant increase of *MSMB* expression. (A) Paired gRNA (rs10993994-g4a and -g4b) mediated CRISPR/Cas9 deletion of rs10993994 region was confirmed by PCR amplification with flanking primer pair rs10993994-F1 and -R1. PCR product runs at 274bp on agarose gel with deletion, and at 479bp without deletion. (B) Real time qPCR showed 2.8 folds *MSMB* over-expression in prostate cancer LNCaP cells with bulk deletion but not in the gastric cancer AGS cells. The expression of downstream *NCOA4* gene is down-regulated. (C) Real time qPCR of single clones generated from above bulk transfection. MSMB over-expression was seen in two (clone 5, 6) out of the three clones (clone 4, 5, 6) with rs10993994 C allele (C/del) but not the T allele (T/del, clone 2, 3, 4).

## Discussion

Here, we have unequivocally demonstrated dramatic allelic effects on *MSMB* expression at two prostate-cancer associated SNPs. Notably, despite these two SNPs being in strong linkage disequilibrium in European populations, each SNP appears to independently influence *MSMB* expression. One can imagine a model in which it is the set of alleles on a haplotype, each exerting an effect, that leads to the observed phenotype rather than the typically assumed model in which there is a single causal allele in LD with other SNPs. Consistent with this hypothesis, reporter assays based on phenotype-associated human haplotypes have demonstrated such additive effects as well[38].

Previous correlative work had implicated both *MSMB* and *NCOA4* as target genes of rs10993994 and potential mediators of the SNP’s effect on prostate cancer risk[25], though studies of protein expression found that rs10993994 is only associated with β-MSP, and not NCOA4, expression[39]. Our results demonstrate that regulatory regions encompassing both rs10993994 and rs7098889 directly affect the levels of *MSMB* only and have no effect on *NCOA4. MSMB* codes for β-MSP, a major secretory product of the prostate. It is widely secreted by multiple mucosal tissues[40] and has been proposed to manifest both fungicidal activity[41] and tumor suppressive properties[25,42,43]. The levels of β-MSP in blood are negatively correlated with risk of a prostate cancer diagnosis[44] and positively associated with recurrence-free survival [45]. These findings are consistent with a direct role for *MSMB* in prostate cancer etiology. However, we cannot exclude a potential role for *NCOA4*, mediated by previously unidentified SNPs in LD with rs10993994, either instead of or in addition to *MSMB*.

The number of GWAS loci for which causal variants and their downstream effect have been identified using genome editing is relatively limited[9,11,13–19,46]. One limiting factor is that in many cases, genome editing revealed that the allelic effect on target gene expression is very mild. In contrast, here we have shown extensive overexpression conferred by altering SNP-containing regulatory regions. Second, the low efficiency of homology-directed repair limits large-scale application of CRISPR/Cas9. Our alternative approach of using a paired gRNA system[34,35] to delete single copies of heterozygous alleles allows us to observe the effect of each variant allele in isolation.

As GWAS have identified numerous genetic polymorphisms associated with increased risk of cancer and other diseases, the next big challenge is to understand how they mediate pathogenesis, especially for regulatory variants. The majority of these variants are found in the non-coding region of the genome[1–3] and enriched in regulatory regions identified by large-scale functional genomics projects including ENCODE, Roadmap Epigenome and FANTOM5[2–6]. These observations together with a recent report that 82% of cancers depend on RNA expression level[20] changes indicate the importance of studying the relationship between disease risk variants and gene expression. Our study for the first time demonstrates dramatic cell-specific and gene-specific effect on gene expression mediated by GWAS-identified risk variants and provides an efficient way for further systematic study of the function of other GWAS variants.

## Materials and methods

### CRISPR/Cas9 mediated genome editing with paired gRNAs

We designed paired guide RNAs flanking either the rs7098889 (rs7098889-g1 and rs7098889-g4, S1 table) or the rs10993994 (rs10993994-g4a and rs10993994-g4b, S2 table) sites with the Broad Institute CRISPR Design tool (crispr.mit.edu). Guide RNAs were chosen based on the best specificity while maintaining a deletion size of around 200bp. Additionally, since rs1099394 is so close to the transcription start site (TSS) of *MSMB*, the downstream flanking gRNA was chosen to preserve the TSS. Each pair of gRNAs were cloned into pSpCas9(BB)-2A-GFP (PX458) (Addgene, Cambridge, MA, Plasmid 48138) and pSpCas9(BB)-2A-Puro (PX459) V2.0 (Addgene, Plasmid 62988) vectors respectively with CRISPR cloning service from Genscript (Piscataway, NJ). The PX458 vector expresses the Cas9 nuclease, upstream gRNA and the EGFP transfection marker, while the PX459 vector expresses the Cas9 nuclease, downstream gRNA and a puromycin selection marker. The combined use of PX458 and PX459 for paired gRNAs transfection provides convenience to both visualize transfection efficiency under a fluorescence microscope and to do post-transfection puromycin drug selection.

### Cell culture and transfection

Both the prostate cancer cell line LNCaP (CRL-1740) and the gastric cancer cell line AGS were obtained from ATCC (ATCC, Rockville, MD). The LNCaP cells were cultured in RPMI 1640 medium (Gibco 11875-093, Life Technologies) supplemented with 15% fetal bovine serum (FBS) and the AGS cells were cultured in F12K medium (Gibco 21127-022, Life Technologies) with 10% FBS, both with the presence of 1% penicillin/streptomycin (Gibco 15140-122, Life Technologies). Cells were kept in standard 37°C, 5% CO2 incubator. Cells were split the day before transfection and allowed to reach 30-70% confluence on the day of transfection. To generate deletion of the rs7098889 site, 2μg each of PX458-rs7098889-g1 and PX459v2-rs7098889-g4 plasmids were mixed with 250μl Opti-MEM (Gibco 31985-070, Life Technologies), then gently mixed with room temperature 10μl Lipofectmine 2000/250μl Opti-MEM mix. After 20 min incubation at room temperature, the mix was added evenly to cells. Cells were put back in the incubator for 4-6 hours before changing to 37°C warm and fresh medium. As control, the empty vector PX458 and PX459v2 pair were transfected in parallel. 36-48 hours later, transfected GFP positive cells were observed under fluorescent microscopy to ensure successful transfection. Puromycin selection was begun by incubating cells in the presence of 2μg/ml puromycin (Santa Cruz Biotechnology sc-108071). Approximately 3-7 days later, puromycin was reduced to 0.5μg/μl for post-selection cell expansion. For single cell cloning, 1μg of each plasmid was used in transfection to reduce transfection and deletion efficiency.

### Genomic PCR and identification of isogenic allelic deletion by Sanger sequencing

Genomic DNA was extracted from bulk transfected LNCaP or AGS cells upon completion of puromycin selection using DNeasy kit (Qiagen, Germany), or from LNCaP single clones derived from the bulk transfections. Deletion was confirmed by PCR amplification using primers flanking the deletion region, rs7098889-For1 and rs7098889-Rev1 for rs7098889 site (S1 Table, synthesized by Invitrogen); and rs10993994-F1 and rs10993994-R1 (S2 Table, synthesized by Invitrogen) for rs10993994 site. Touchdown PCR was used (95°C 1 min; 10 cycles of 95°C 15 sec, 68°C −1°C/cycle 15 sec, 72°C 30 sec; 25 cycles of 95°C 15 sec, 60°C 15 sec, 72°C 30 sec; 72°C 5 min, 4°C incubate). PCR products were resolved on 1.5% agarose gel stained by SYBR green and visualized under UV light. Alleles with deletion end up with 178bp band compared to the 369bp no deletion band for rs7098889 deletion; 274bp vs. 479bp for rs10993994 deletion. All bands were excised from the agarose gel, purified by QIAquick Gel Extraction Kit (Qiagen, Germany), and sent for Sanger sequencing (Genewiz) for sequence validation. Only the clones with correct deletion junction and correct wild type sequence were used for further analysis.

### Total RNA extraction and real time qPCR analysis

To compare gene expression changes of *MSMB, NCOA4, KLK2* and *KLK3* from different transfection and isogenic clones, cells were harvested at 60-90% confluence for total RNA extraction with Qiagen RNeasy Mini kit (Qiagen, Hilden, Germany) and quantified by Nanodrop spectrophotometer (ThermoScientific, ND-8000). 1μg extracted RNA were then reverse transcribed into cDNA with the High Capacity cDNA Reverse Transcription Kit (Applied Biosystems #4368814). For real time PCR, TaqMan gene-specific primers were ordered from Life Technologies for *MSMB* (Hs00159303_m1), *NCOA4* (Hs01033772_g1), *KLK2* (Hs00428383_m1) and *KLK3* (Hs02576345). GAPDH was used as internal control. qPCR reactions were setup according to the TaqMan Gene Expression Assays protocol and performed on a ViiA7 real time PCR system (Applied Biosystems, Life Technologies). Each sample was amplified in duplicate, average gene expression and standard deviation were calculated. Relative gene expression was analyzed with the ΔΔC_T_ method (Applied Biosystems, cms_042380).

### Western blot

Cells from six-well plates were collected by cold PBS, whole cell lysate were prepared using cold NP40 buffer (150mM NaCl, 1% Igepal CA630 and 50mM Tris-HCl, pH 8.0) supplemented with Protease Arrest™ (Calbiochem #KP14001, 1:50 dilution). After incubation on ice for 30 minutes and centrifugation for 10 minutes at 12,000 rpm, supernatant was recovered and protein concentrations were measured using the BioRad Protein Assay Dye Reagent (BioRad 500-0006). 50μg protein was pipetted into each lane and separated on a SDS-PAGE gel (Bolt 4-12% Bis-Tris Plus, Invitrogen, Life Technologies) and the Precision Plus Protein Standards (BioRad) was used as molecular weight marker. Transfer was done on Bio-Rad Trans-Blot Turbo Transfer System onto 0.2 um PVDF membrane (BioRad #1704156). After blocking in 5% milk at room temperature for 1 hour, membranes were incubated in primary antibody diluted in TBST buffer with 1% BSA (Sigma #A9647) for overnight at 4°C. Followed by 3 times 10 minutes wash in TBST buffer (10X Bio-Rad TBS plus 0.05% Tween-20), membranes were incubated another hour at room temperature in secondary antibody diluted in 2.5% milk TBST buffer. After final wash, the results were visualized by the Bio-Rad ClarityTM Western ECL substrate (Bio-Rad #170-5061). Antibodies used are: anti-β-MSP (Origene clone 6C7 #TA501072, 1:3000, lot #A01); anti-β-actin (Sigma clone AC-15, #A5441, 1:5000, lot #122M4782); anti-mouse IgG-HRP (ThermoScientific Pierce Antibody #31432, 1:5000).

### Immunodetection of β-MSP, hK2 and PSA secretion

All immunoassay measurements of PSA, hK2, and MSP were conducted blinded on the Victor instrument (Perkin-Elmer, Turku, Finland) using the dual-label DELFIA Prostatus^®^total/free PSA-Assay (Perkin-Elmer, Turku, Finland) calibrated against the World Health Organization (WHO) 96/670 (PSA-WHO) and WHO 68/668 (free PSA-WHO) standards. Production and purification of the polyclonal rabbit anti-MSP antibody, protocols for biotinylation and Europium labeling of the anti-MSP antibody, and performance of the MSP-immunoassay were performed as previously reported [36]. Duplicate samples were read, average and standard deviation were plotted.

### Single cell cloning

Upon completion of puromycin selection, bulk PX458-rs7098889-g1 and PX459v2-rs7098889-g4 transfected LNCaP cells or the bulk PX458-rs10993994-g4a and PX459v2-rs10993994-g4b transfected LNCaP cells were plated into 96-well plates (Falcon) by serial dilution. Cells were cultured in LNCaP condition medium filtered by 0.22μm Millex membrane (Millipore) until colonies starting form in about 3 weeks. Single clones were then trypsinized and transferred to 24-well plates followed by 6 well plates in triplicates, and used for genomic DNA extraction followed by PCR and Sanger sequencing genotype identification, freezing clones, and future experiments, respectively.

### Allelic expression analysis

Total RNA was extracted from bulk transfected LNCaP and validated single clones, followed by reverse transcription (High Capacity cDNA Reverse Transcription Kit, Applied Biosystems #4368814), cDNA then amplified by PCR reaction with primers flanking *MSMB* 360A/T site. Agarose gel purified PCR products were then subjected to Sanger sequencing by Genewiz and returned chromatogram sequence were visualized in 4Peaks.

## Supporting information

**S1Fig. ELISA assay in LNCaP cells with CRISPR/Cas9 mediated rs7098889 deletion.**

**S2 Fig. Real time qPCR of *NCOA4, KLK2* and *KLK3* in rs7098889 bulk deletion and single clones.**

**S3 Fig. CRISPR/Cas9 mediated deletion of rs7098889 or rs10993994 in RWPE-1 cells.**

**S4 Fig. Real time qPCR of *NCOA4, KLK2* and *KLK3* in rs10993994 single clones.**

**S1 Table. Guide RNA and primer sequence for rs7098889 site.**

**S2 Table. Guide RNA and primer sequence for rs10993994 site.**

**S1 File. Sanger sequencing alignment of LNCaP rs7098889 single clones.**

**S2 File. Sequence alignment of LNCaP rs10993994 single clones.**

## Data availability

Cell clones generated in the study are available upon request. Contact: Robert J Klein

## Author contributions

X.W. designed and conducted this research; J.E.H. performed the analysis of FANTOM5 data; X.X. identified the heterozygous coding variant; X.G. assisted with experiments; D.M. performed the immunoassays; H.G.L. supervised the immunoassay measurements; R.J.K. designed and supervised this research.

## References

1. Manolio TA. Genomewide Association Studies and Assessment of the Risk of Disease. Feero WG, Guttmacher AE, editors. N Engl J Med. 2010;363: 166–176. doi:10.1056/NEJMra0905980

2. Maurano MT, Humbert R, Rynes E, Thurman RE, Haugen E, Wang H, et al. Systematic localization of common disease-associated variation in regulatory DNA. Science. 2012;337: 1190–5. doi:10.1126/science.1222794

3. Dunham I, Kundaje A, Aldred SF, Collins PJ, Davis CA, Doyle F, et al. An integrated encyclopedia of DNA elements in the human genome. Nature. 2012;489: 57–74. doi:10.1038/nature11247

4. Kundaje A, Meuleman W, Ernst J, Bilenky M, Yen A, Heravi-Moussavi A, et al. Integrative analysis of 111 reference human epigenomes. Nature. 2015;518: 317–330. doi:10.1038/nature14248

5. Andersson R, Gebhard C, Miguel-Escalada I, Hoof I, Bornholdt J, Boyd M, et al. An atlas of active enhancers across human cell types and tissues. Nature. 2014;507: 455–461. doi:10.1038/nature12787

6. Price AL, Spencer CCA, Donnelly P. Progress and promise in understanding the genetic basis of common diseases. Proceedings Biol Sci. The Royal Society; 2015;282: 20151684. doi:10.1098/rspb.2015.1684

7. Ardlie KG, Deluca DS, Segre A V., Sullivan TJ, Young TR, Gelfand ET, et al. The Genotype-Tissue Expression (GTEx) pilot analysis: Multitissue gene regulation in humans. Science (80-). 2015;348: 648–660. doi:10.1126/science.1262110

8. Levenstien MA, Klein RJ. Predicting functionally important SNP classes based on negative selection. BMC Bioinformatics. 2011;12: 26. doi:10.1186/1471-2105-12-26

9. Yao L, Tak YG, Berman BP, Farnham PJ. Functional annotation of colon cancer risk SNPs. Nat Commun. 2014;5: 5114. doi:10.1038/ncomms6114

10. Claussnitzer M, Dankel SN, Kim K-H, Quon G, Meuleman W, Haugen C, et al. FTO Obesity Variant Circuitry and Adipocyte Browning in Humans. N Engl J Med. 2015;373: 895–907. doi:10.1056/NEJMoa1502214

11. Spisak S, Lawrenson K, Fu Y, Csabai I, Cottman RT, Seo JH, et al. CAUSEL: an epigenome- and genome-editing pipeline for establishing function of noncoding GWAS variants. Nat Med. 2015;21: 1357–1363. doi:10.1038/nm.3975

12. Tak YG, Hung Y, Yao L, Grimmer MR, Do A, Bhakta MS, et al. Effects on the transcriptome upon deletion of a distal element cannot be predicted by the size of the H3K27Ac peak in human cells. Nucleic Acids Res. 2016;44: 4123–4133. doi:10.1093/nar/gkv1530

13. Won H, de la Torre-Ubieta L, Stein JL, Parikshak NN, Huang J, Opland CK, et al. Chromosome conformation elucidates regulatory relationships in developing human brain. Nature. 2016;538: 523–527. doi:10.1038/nature19847

14. Jin H-J, Jung S, DebRoy AR, Davuluri R V. Identification and validation of regulatory SNPs that modulate transcription factor chromatin binding and gene expression in prostate cancer. Oncotarget. 2016;7: 54616–54626. doi:10.18632/oncotarget.10520

15. Fulco CP, Munschauer M, Anyoha R, Munson G, Grossman SR, Perez EM, et al. Systematic mapping of functional enhancer-promoter connections with CRISPR interference. Science. 2016;354: 769–773. doi:10.1126/science.aag2445

16. Liu H, Leslie EJ, Carlson JC, Beaty TH, Marazita ML, Lidral AC, et al. Identification of common non-coding variants at 1p22 that are functional for non-syndromic orofacial clefting. Nat Commun. 2017;8: 14759. doi:10.1038/ncomms14759

17. Soldner F, Stelzer Y, Shivalila CS, Abraham BJ, Latourelle JC, Barrasa MI, et al. Parkinson-associated risk variant in distal enhancer of α-synuclein modulates target gene expression. Nature. 2016;533: 95–99. doi:10.1038/nature17939

18. Maharry SE, Walker CJ, Liyanarachchi S, Mehta S, Patel M, Bainazar MA, et al. Dissection of the Major Hematopoietic Quantitative Trait Locus in Chromosome 6q23.3 Identifies miR-3662 as a Player in Hematopoiesis and Acute Myeloid Leukemia. Cancer Discov. 2016;6: 1036–1051. doi:10.1158/2159-8290.CD-16-0023

19. Grampp S, Platt JL, Lauer V, Salama R, Kranz F, Neumann VK, et al. Genetic variation at the 8q24.21 renal cancer susceptibility locus affects HIF binding to a MYC enhancer. Nat Commun. 2016;7: 13183. doi:10.1038/ncomms13183

20. Tsherniak A, Vazquez F, Montgomery PG, Weir BA, Kryukov G, Cowley GS, et al. Defining a Cancer Dependency Map. Cell. 2017;170: 564–576.e16. doi:10.1016/j.cell.2017.06.010

21. Grönberg H, Damber L, Damber JE. Studies of genetic factors in prostate cancer in a twin population. J Urol. 1994;152: 1484-7-9. Available: http://www.ncbi.nlm.nih.gov/pubmed/7933190

22. Lichtenstein P, Holm N V., Verkasalo PK, Iliadou A, Kaprio J, Koskenvuo M, et al. Environmental and Heritable Factors in the Causation of Cancer — Analyses of Cohorts of Twins from Sweden, Denmark, and Finland. N Engl J Med. 2000;343: 78–85. doi:10.1056/NEJM200007133430201

23. Mucci LA, Hjelmborg JB, Harris JR, Czene K, Havelick DJ, Scheike T, et al. Familial Risk and Heritability of Cancer Among Twins in Nordic Countries. JAMA. NIH Public Access; 2016;315: 68–76. doi:10.1001/jama.2015.17703

24. Al Olama AA, Kote-Jarai Z, Berndt SI, Conti D V, Schumacher F, Han Y, et al. A meta-analysis of 87,040 individuals identifies 23 new susceptibility loci for prostate cancer. Nat Genet. 2014;46: 1103–1109. doi:10.1038/ng.3094

25. Pomerantz MM, Shrestha Y, Flavin RJ, Regan MM, Penney KL, Mucci LA, et al. Analysis of the 10q11 Cancer Risk Locus Implicates MSMB and NCOA4 in Human Prostate Tumorigenesis. Ford JM, editor. PLos Genet. 2010;6: e1001204. doi:10.1371/journal.pgen.1001204

26. Grisanzio C, Werner L, Takeda D, Awoyemi BC, Pomerantz MM, Yamada H, et al. Genetic and functional analyses implicate the NUDT11, HNF1B, and SLC22A3 genes in prostate cancer pathogenesis. Proc Natl Acad Sci U S A. 2012;109: 11252–11257. doi:10.1073/pnas.1200853109

27. Penney KL, Sinnott JA, Tyekucheva S, Gerke T, Shui IM, Kraft P, et al. Association of prostate cancer risk variants with gene expression in normal and tumor tissue. Cancer Epidemiol Biomarkers Prev. 2015;24: 255–260. doi:10.1158/1055-9965.EPI-14-0694-T

28. Xu X, Hussain WM, Vijai J, Offit K, Rubin MA, Demichelis F, et al. Variants at IRX4 as prostate cancer expression quantitative trait loci. Eur J Hum Genet. 2014;22: 558–563. doi:10.1038/ejhg.2013.195

29. Savblom C, Hallden C, Cronin AM, Sall T, Savage C, Vertosick EA, et al. Genetic variation in KLK2 and KLK3 is associated with concentrations of hK2 and PSA in serum and seminal plasma in young men. Clin Chem. 2014;60: 490–499. doi:10.1373/clinchem.2013.211219

30. Xu X, Valtonen-André C, Sävblom C, Halldén C, Lilja H, Klein RJ. Polymorphisms at the microseminoprotein-?? locus associated with physiologic variation in_??-microseminoprotein and prostate-specific antigen levels. Cancer Epidemiol Biomarkers Prev. 2010;19: 2035–2042. doi:10.1158/1055-9965.EPI-10-0431

31. Gusev A, Shi H, Kichaev G, Pomerantz M, Li F, Long HW, et al. Atlas of prostate cancer heritability in European and African-American men pinpoints tissue-specific regulation. Nat Commun. 2016;7: 10979. doi:10.1038/ncomms10979

32. Whitington T, Gao P, Song W, Ross-Adams H, Lamb AD, Yang Y, et al. Gene regulatory mechanisms underpinning prostate cancer susceptibility. Nat Genet. 2016;48: 387–397. doi:10.1038/ng.3523

33. Lou H, Yeager M, Li H, Bosquet JG, Hayes RB, Orr N, et al. Fine mapping and functional analysis of a common variant in MSMB on chromosome 10q11.2 associated with prostate cancer susceptibility. Proc Natl Acad Sci. 2009;106: 7933–7938. doi:10.1073/pnas.0902104106

34. Ran FA, Hsu PD, Wright J, Agarwala V, Scott DA, Zhang F. Genome engineering using the CRISPR-Cas9 system. Nat Protoc. 2013;8: 2281–2308. doi:10.1038/nprot.2013.143

35. Zheng Q, Cai X, Tan MH, Schaffert S, Arnold CP, Gong X, et al. Precise gene deletion and replacement using the CRISPR/Cas9 system in human cells. Biotechniques. 2014;57: 115–24. doi:10.2144/000114196

36. Haiman CA, Stram DO, Vickers AJ, Wilkens LR, Braun K, Valtonen-André C, et al. Levels of beta-microseminoprotein in blood and risk of prostate cancer in multiple populations. J Natl Cancer Inst. 2013;105: 237–43. doi:10.1093/jnci/djs486

37. Chang B-L, Cramer SD, Wiklund F, Isaacs SD, Stevens VL, Sun J, et al. Fine mapping association study and functional analysis implicate a SNP in MSMB at 10q11 as a causal variant for prostate cancer risk. Hum Mol Genet. 2009;18: 1368–1375. doi:10.1093/hmg/ddp035

38. Guo C, Ludvik AE, Arlotto ME, Hayes MG, Armstrong LL, Scholtens DM, et al. Coordinated regulatory variation associated with gestational hyperglycaemia regulates expression of the novel hexokinase HKDC1. Nat Commun. 2015;6: 6069. doi:10.1038/ncomms7069

39. FitzGerald LM, Zhang X, Kolb S, Kwon EM, Liew YC, Hurtado-Coll A, et al. Investigation of the Relationship Between Prostate Cancer and MSMB and NCOA4 Genetic Variants and Protein Expression. Hum Mutat. 2013;34: 149–156. doi:10.1002/humu.22176

40. Weiber H, Andersson C, Murne A, Rannevik G, Lindström C, Lilja H, et al. Beta microseminoprotein is not a prostate-specific protein. Its identification in mucous glands and secretions. Am J Pathol. 1990;137: 593–603. Available: http://www.ncbi.nlm.nih.gov/pubmed/2205099

41. Edström Hägerwall AML, Rydengård V, Fernlund P, Mörgelin M, Baumgarten M, Cole AM, et al. β-Microseminoprotein Endows Post Coital Seminal Plasma with Potent Candidacidal Activity by a Calcium- and pH-Dependent Mechanism. Feldmesser M, editor. Plos Pathog. 2012;8: e1002625. doi:10.1371/journal.ppat.1002625

42. Garde S, Sheth A, Porter AT, Pienta KJ. Effect of prostatic inhibin peptide (PIP) on prostate cancer cell growth in vitro and in vivo. Prostate. 1993;22: 225–33. Available: http://www.ncbi.nlm.nih.gov/pubmed/8488155

43. Lokeshwar BL, Hurkadli KS, Sheth AR, Block NL. Human prostatic inhibin suppresses tumor growth and inhibits clonogenic cell survival of a model prostatic adenocarcinoma, the Dunning R3327G rat tumor. Cancer Res. 1993;53: 4855–9. Available: http://www.ncbi.nlm.nih.gov/pubmed/8402673

44. Whitaker HC, Kote-Jarai Z, Ross-Adams H, Warren AY, Burge J, George A, et al. The rs10993994 risk allele for prostate cancer results in clinically relevant changes in microseminoprotein-beta expression in tissue and urine. PLos One. 2010;5: e13363. doi:10.1371/journal.pone.0013363

45. Bjartell AS, Al-Ahmadie H, Serio AM, Eastham JA, Eggener SE, Fine SW, et al. Association of Cysteine-Rich Secretory Protein 3 and -Microseminoprotein with Outcome after Radical Prostatectomy. Clin Cancer Res. 2007;13: 4130–4138. doi:10.1158/1078-0432.CCR-06-3031

46. Claussnitzer M, Dankel SN, Kim K-H, Quon G, Meuleman W, Haugen C, et al. FTO Obesity Variant Circuitry and Adipocyte Browning in Humans. N Engl J Med. 2015;373: 895–907. doi:10.1056/NEJMoa1502214

